# A fully-automated, robust, and versatile algorithm for long-term budding yeast segmentation and tracking

**DOI:** 10.1101/444018

**Authors:** N Ezgi Wood, Andreas Doncic

## Abstract

Live cell time-lapse microscopy, a widely-used technique to study gene expression and protein dynamics in single cells, relies on segmentation and tracking of individual cells for data generation. The potential of the data that can be extracted from this technique is limited by the inability to accurately segment a large number of cells from such microscopy images and track them over long periods of time. Existing segmentation and tracking algorithms either require additional dyes or markers specific to segmentation or they are highly specific to one imaging condition and cell morphology and/or necessitate manual correction. Here we introduce a fully automated, fast and robust segmentation and tracking algorithm for budding yeast that overcomes these limitations. Full automatization is achieved through a novel automated seeding method, which first generates coarse seeds, then automatically fine-tunes cell boundaries using these seeds and automatically corrects segmentation mistakes. Our algorithm can accurately segment and track individual yeast cells without any specific dye or biomarker. Moreover, we show how existing channels devoted to a biological process of interest can be used to improve the segmentation. The algorithm is versatile in that it accurately segments not only cycling cells with smooth elliptical shapes, but also cells with arbitrary morphologies (e.g. sporulating and pheromone treated cells). In addition, the algorithm is largely independent of the specific imaging method (bright-field/phase) and objective used (40X/63X). We validate our algorithm’s performance on 9 cases each entailing a different imaging condition, objective magnification and/or cell morphology. Taken together, our algorithm presents a powerful segmentation and tracking tool that can be adapted to numerous budding yeast single-cell studies.

## Introduction

Traditional life science methods that rely on the synchronization and homogenization of cell populations have been used with great success to address numerous questions; however, they mask dynamic cellular events such as oscillations, all-or-none switches, and bistable states [1-5]. To capture and study such behaviors, the process of interest should be followed over time at single cell resolution [6-8]. A widely used method to achieve this spatial and temporal resolution is live-cell time-lapse microscopy [9], which has two general requirements for extracting single-cell data: First, single-cell boundaries have to be identified for each time-point (segmentation), and second, cells have to be tracked over time (tracking) [10, 11].

One of the widely-used model organisms in live-cell microscopy is budding yeast *Sacchromyces cerevisiae*, which is easy to handle, has tractable genetics, and a short generation time [12, 13]. Most importantly in the context of image analysis, budding yeast cells have smooth cell boundaries and are mostly stationary while growing, which can be exploited by segmentation and tracking algorithms. Thus, in contrast to many mammalian segmentation approaches that segment only the nucleus, use dyes to stain the cytoplasm [14-17], use manual cell tracking [18] or extract features using segmentation-free approaches [19], we expect yeast segmentation to be completely accurate using only phase or bright-field images. Hence, budding yeast segmentation and tracking pose a complex optimization problem in which we strive to simultaneously achieve automation, accuracy, and general applicability with no or limited use of biomarkers.

Several different methods and algorithms have been created to segment and track yeast cells. To reach high accuracy, some of these algorithms rely on images where cell boundaries and/or the cell nuclei are stained [20-22]. However, with staining, one or several fluorescent channels are ‘occupied’, which limits the number of available channels that could be used to collect information about cellular processes [23]. In addition, using fluorescent light for segmentation increases the risk for photo-toxicity and bleaching [24]. Thus, it is desirable to segment and track cells using only bright-field or phase images.

Another commonly used method, ‘2D active contours’, fits parametrized curves to cell boundaries [25]. Existing yeast segmentation algorithms using this method typically take advantage of the elliptical shape of cycling yeast cells [26-28]. Another way to take advantage of the prior information on cell shape is to create a shape library where shapes from an ellipse library and cells are matched [29]. Although these methods can be very accurate, they tend to be computationally expensive [29], and, to the best of our knowledge, they are not tested on any non-ellipsoidal morphologies, e.g. sporulating or pheromone treated cells. Moreover, in many cases they have to be fine-tuned to the specific experimental setup used [27, 29].

Here we present a fully automated segmentation and tracking algorithm for budding yeast cells. The algorithm builds on our previously published algorithm [30], improves its accuracy and speed significantly, and fully automatizes it by introducing a novel automated seeding step. This seeding step incorporates a new way for automated cell boundary fine-tuning and automated correction of segmentation errors. Our algorithm is parallelizable, and thus fast, and segments arbitrary cell shapes with high accuracy. Our algorithm does not rely on segmentation specific staining or markers. Still, we show how information about cell locations can be incorporated into the segmentation algorithm using fluorescent channels that are *not* devoted to segmentation. To demonstrate the versatility of our algorithm we validate it on 9 different example cases each with a different cell morphology, objective magnification and/or imaging method (phase / bright-field).

## Results

### Automated seeding

When segmenting yeast cells over time, it is advantageous to start at the last time-point and segment the images backwards in time [30], because all cells are present at the last time point due to the immobility of yeast cells. Thus, instead of attempting the harder problem of detecting newborn cells (buds), we only have to follow existing cells backwards in time until they are born (disappear). To segment the cells, we therefore need an initial segmentation of the last time-point, which is fed to the main algorithm that uses the segmentation of the previous time point as the seed for the next time point.

This seeding step was previously a bottleneck since it was semi-automated and required user-input. To fully automate the segmentation algorithm, we developed a novel method to automate this seeding step. Here we present the general outline of this method. For a detailed explanation see the supplementary material and the accompanying annotated software.

The automated seeding algorithm has two main steps (Fig 1): First, watershed algorithm is applied to the pre-processed image of the last time point (Fig 1A-C). Second, the resulting watershed lines are automatically fine-tuned, and segmentation mistakes are automatically corrected (Fig 1D and 1E).

**Fig 1.**
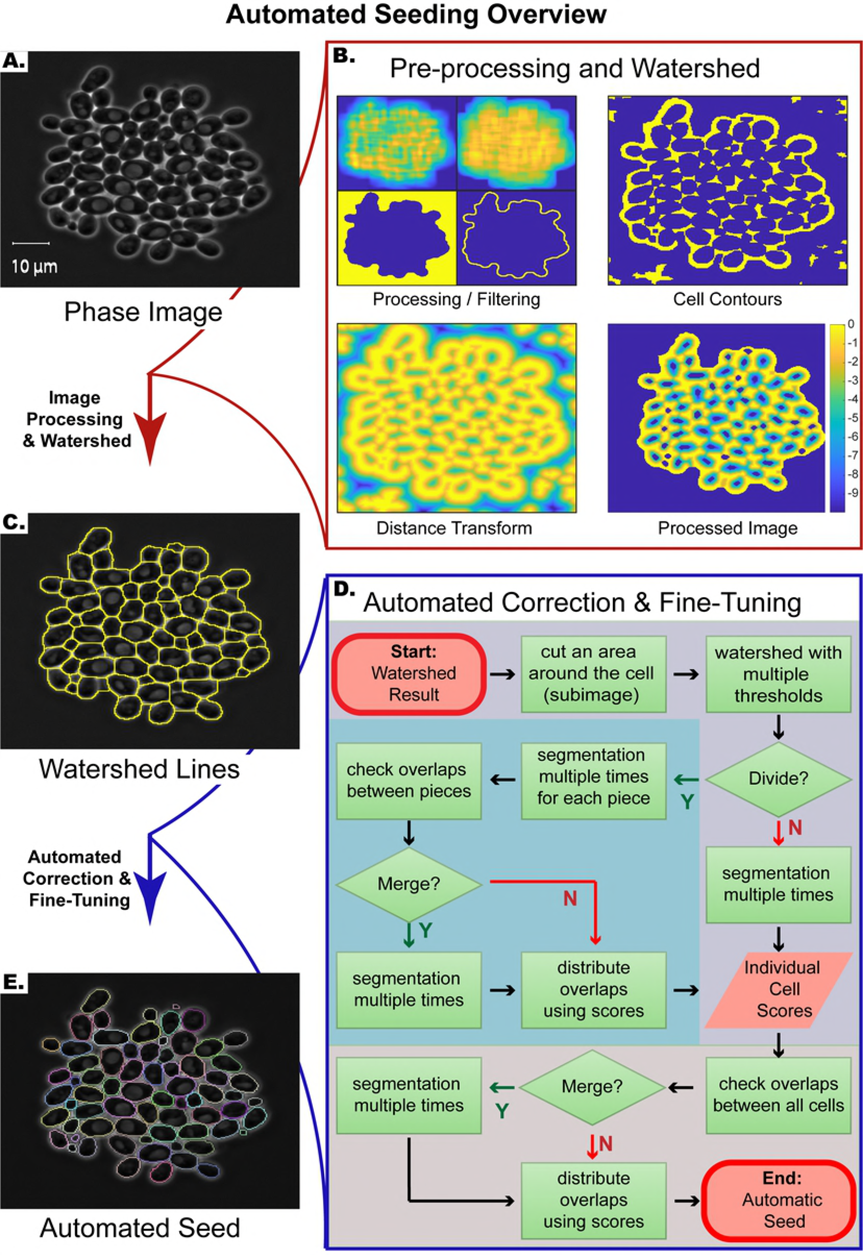
Automated seeding overview. **(A)** Example phase image. **(B)** First step of automated seeding algorithm: Pre-processing and watershed. In this step, the watershed transform is applied to the processed image. **(C)** Phase image with watershed lines (yellow). **(D)** Flowchart of the second step of automated seeding: Automated correction and fine-tuning. At this step, the cell boundaries are automatically fine-tuned, and segmentation errors are automatically corrected. **(E)** The result of the automated seeding step. Each cell boundary is marked with a different color.

### Pre-processing and watershed

During this step, the image is processed before the application of the watershed transform, with the aim of getting only one local minimum at each cell interior, so that each cell area will be associated with one segmented region after the application of the watershed transform. To this end, the image is first coarsely segmented to determine the cell and non-cell (background) regions of the image (Fig 1B, Processing/Filtering, binary image on the bottom left). Based on this coarse segmentation, we only focus on the cell colonies. Next, cell contours and interstices are identified by exploiting the fact that they are brighter than the background pixels and cell interiors (Fig 1B, Cell Contours). To detect such pixels, we use mean and standard deviation filtering (Fig 1B, Processing/Filtering, top images) and label pixels that are brighter than their surroundings as cell contour pixels. Once these cell contour pixels are determined, we apply a distance transform to this binary image and further process the transformed image (Fig1B, Distance Transform and Processed Image). Next, we apply a watershed transform to the resulting image (Fig 1C). Note that even though the watershed lines will separate the cells, they do not mark the exact boundaries (Fig 2A). In addition, sometimes multiple, or lack of, local minima within cells leads to situations where multiple cells are merged as one or a cell is divided into multiple regions (under/over-segmentation, Fig 2B and 2C).

**Fig 2.**
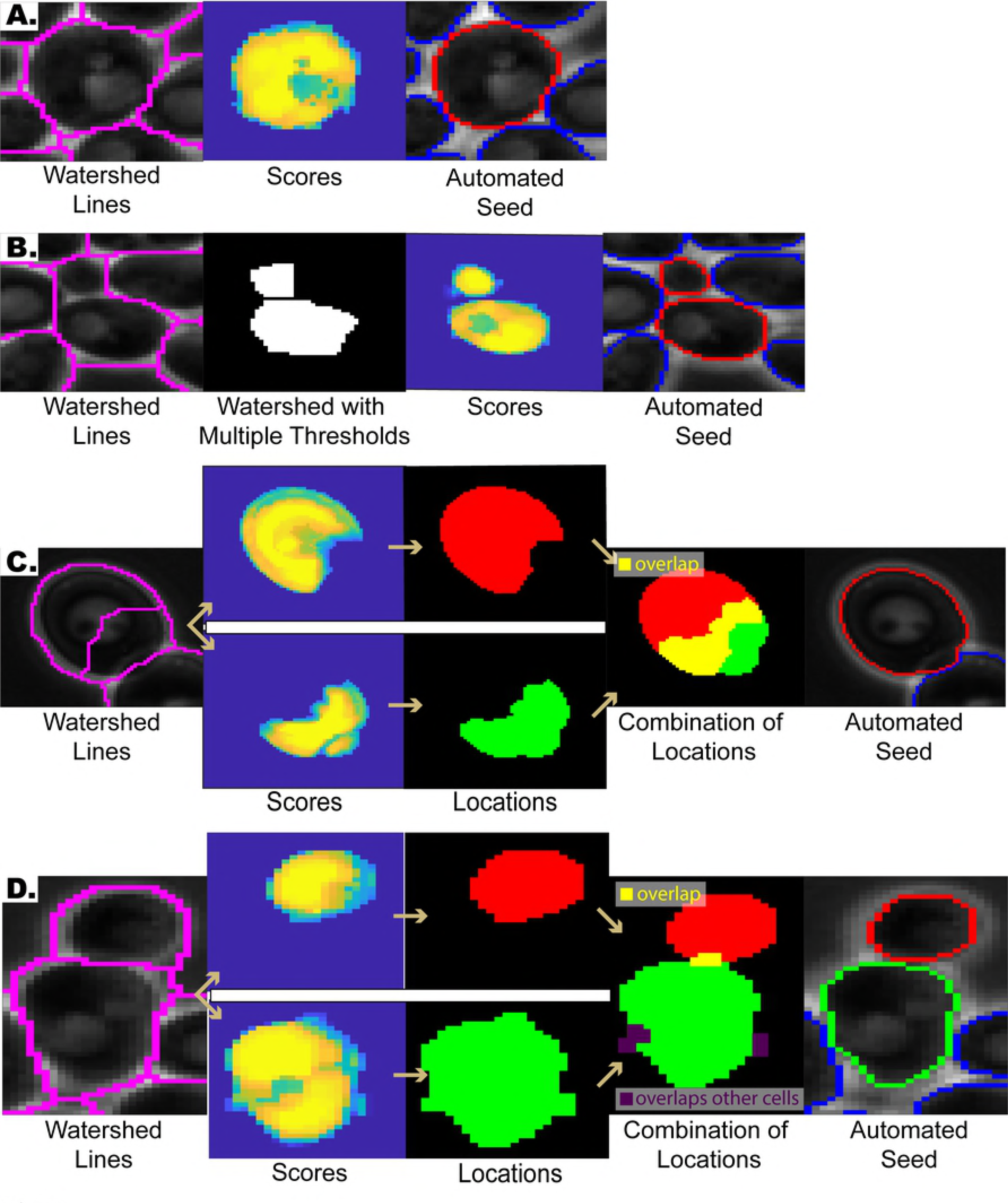
Automated correction & fine-tuning step examples. **(A)** Refining cell boundaries: The watershed lines do not mark the exact cell boundaries (first column, magenta). Our algorithm automatically fine-tunes these watershed lines and marks the correct cell boundary (third column, red). **(B)** Under-segmentation correction: Sometimes the watershed lines merge multiple cells (first column, magenta). Such mistakes are detected and corrected automatically (fourth column, red). **(C)** Over-segmentation correction: Sometimes the watershed lines divide a cell into multiple pieces (first column). After applying the segmentation subroutine several times, each piece converges towards the correct cell segmentation and thus the pieces overlap significantly (fourth column). If the overlap between two pieces are above a certain threshold, then they are merged (fifth column, red). **(D)** Distribution of Overlaps: The algorithm sometimes assigns the same pixels to adjacent cells (Also see section *Distribution of overlapping cell locations*), which leads to overlapping cell locations. Such overlaps (fourth column, yellow) are distributed among the cells based on their scores.

To test our automated seeding step, we applied it to a wide range of example cases: (1) cycling cells imaged by phase contrast with 40X objective and (2) 63X objective, (3) sporulating cells imaged by phase contrast with 40X objective, (4) *cln1 cln2 cln3* cells imaged by phase contrast with 63X objective, (5-8) *cln1 cln2 cln3* cells exposed to 3, 6, 9 and 12nM mating pheromone (α-factor) imaged by phase contrast with 63X objective, and (9) bright-field images of cycling cells imaged with 40X objective. Note that bright-field images were briefly processed before feeding them into the seeding algorithm (see section *Bright-field images*.).

Next, the segmentations were scored manually (Table 1). Cells whose area were correctly segmented over 95% were scored as ‘correct’. A significant fraction of the segmentation mistakes was minor, and they were automatically corrected within 10 time points after the seed was fed into the segmentation and tracking algorithm (Table1). Note that most of the seeding errors emerged from cells with ambiguous cell boundaries, such as dead cells.

### Automated correction and fine-tuning

To refine the cell boundaries and to automatically correct segmentation mistakes, we implemented the second step (Fig 1D), which takes as the input the watershed result from the previous step and gives as the output the final automated seed (Fig 1E). For each cell, this algorithm focuses on a subimage containing the putative cell region determined by the watershed lines. First, the algorithm checks whether the putative cell area contains more than one cell (under-segmentation), i.e. whether the putative cell region needs to be divided. This is achieved by testing the stability of the putative cell location under different parameters: the previous pre-processing and watershed step is applied on the subimage, but this time with multiple thresholds for determining the cell contour pixels. Each threshold has a ‘vote’ for assigning a pixel as a cell pixel or a non-cell pixel, which eventually determines whether the area will be divided. If the putative cell is divided, then each piece is treated separately as an independent cell (Fig1D, blue box). Next, the subimage is segmented using a version of the previously published segmentation subroutine [30] (See supplementary material section *Review of the previously published subroutine*.), in which the image is segmented multiple rounds using the result of the previous segmentation as the seed for the next segmentation. Through these segmentation iterations the coarse seed obtained by the watershed transform converges onto the correct cell boundaries fine-tuning the segmentation. Also, this step generates a *score* for each putative cell, which is an image carrying weights representing how likely each pixel belongs to the cell. These scores are used in case the same pixels are assigned to adjacent cells, leading to overlapping cell locations. If these overlaps are small, the algorithm distributes them among the cells based on the scores generated at the segmentation step (Fig 2D. See also section *Distribution of overlapping cell locations.*). If the intersection between two putative cell regions is above a certain threshold, then the algorithm merges these two regions to correct over-segmentation mistakes (Fig 2C).

**Table 1:** Automated Seeding Performance.

|  | **Initial fraction of**<br>**correctly**<br>**segmented cells**<br>**%** | **Final fraction**<br>**of correctly**<br>**segmented**<br>**cells after 10**<br>**time points** | **Average #**<br>**of time**<br>**points**<br>**needed for**<br>**correction** | **N cells** | **#**<br>**Fields**<br>**of**<br>**View** |
| --- | --- | --- | --- | --- | --- |
| **40X – cycling** | 95.9% | 98.6% | 3.6 | 435 | 2 |
| **63X – cycling** | 95.2% | 97.3% | 5.8 | 293 | 3 |
| **40X- sporulating** | 96.6% | 96.9% | 2 | 352 | 2 |
| **63X – 0 nM** | 92.5% | - | - | 67 | 3 |
| **63X– 3nM** | 93.7% | 95.8% | 5.3 | 143 | 4 |
| **63X – 6 nM** | 90.2% | - | - | 102 | 4 |
| **63X – 9 nM** | 86.24% | 89.9% | 5.2 | 109 | 6 |
| **63X –12nM** | 71.64% | - | - | 67 | 5 |
| **Bright-field** | 95.5% | - | - | 308 | 2 |

0, 3, 6, 9, 12 nM refer to α-factor concentrations used for treating *cln1cln2cln3* cells. #:Number

Finally, we implemented a correction step after the automatic seeding, where faulty seeds can be adjusted or removed semi-automatically. For screening or large-scale applications this step can be omitted with little loss of accuracy.

### Computational Performance

When segmenting an image, the algorithm first segments each cell independent of other cells by focusing on a subimage containing a neighborhood around the cell’s seed. Through parallelization of this step, we significantly improved the speed of our algorithm.

To demonstrate the gain in runtime we segmented an example time-series of images sequentially without parallelization and in parallel with varying number of workers (i.e. parallel processors). The example time-series had 200 images and 360 cells on the last image, which amounted to 25377 segmentation events. With 40 workers the algorithm runs about15-times faster (263 min vs 17 min, Fig 3A). Note that after about 26 workers, there is no significant difference in runtime, since the time gain is limited by the longest serial job. Also, overhead communication time increases with increasing number of workers offsetting the time gain.

**Fig 3.**
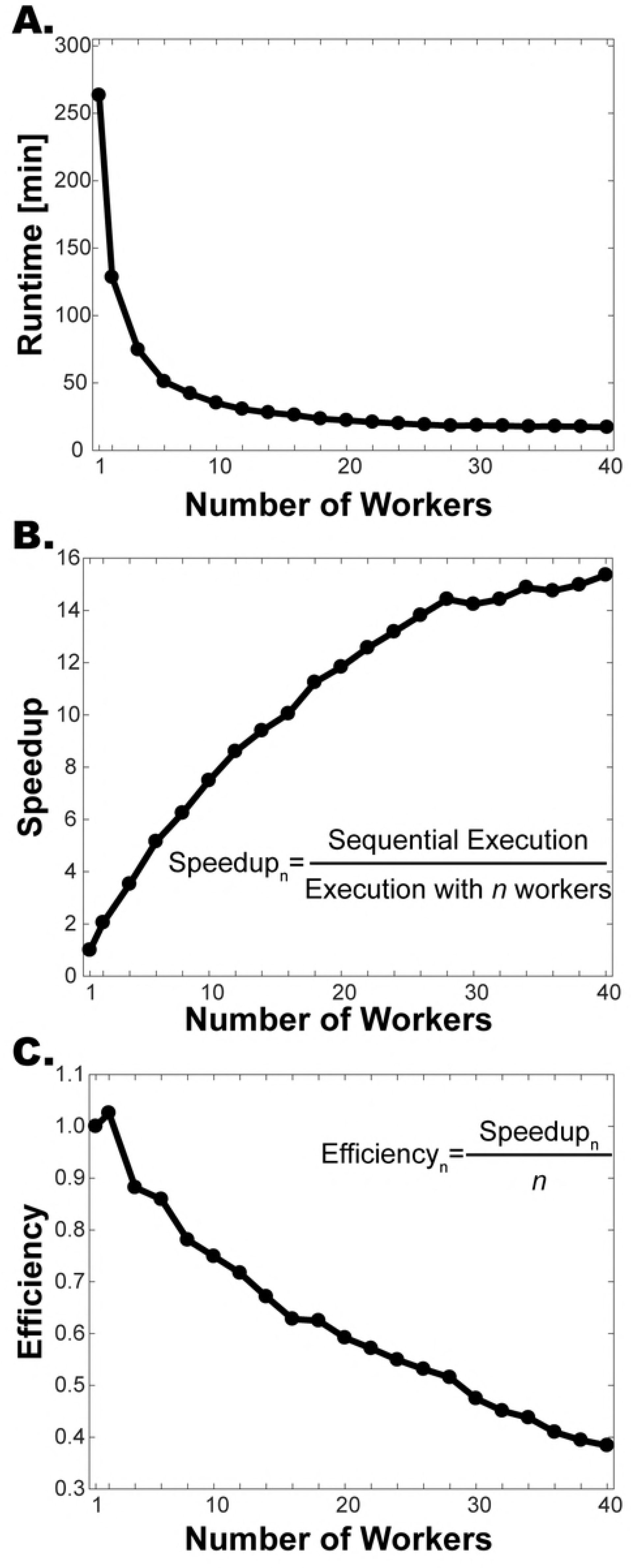
Time gain, speedup and efficiency achieved by parallelization. An example field of view imaged over 10 hours (200 time points, 360 cells at the last time point) was segmented sequentially and in parallel with varying number of workers. **(A)** Runtimes. **(B)** Speedup is calculated by dividing the sequential execution time by the parallel execution time. With 40 workers the algorithm runs 15.4 times faster. **(C)** Efficiency is the speedup per processor. Note that the efficiency goes down as the number of processors increases.

We also calculated the performance measures *speedup* and *efficiency* [31]. The *speedup* is the ratio of the runtime without parallelization to runtime with *n* processors. The speedup increases as the number of workers increases, but eventually levels off (Fig 3B). Next, we calculated the *efficiency*, which is the speedup divided by the number of processors. This gives a measure of how much each processor is used on average [31]. The efficiency is highest for 2 processors and it decreases as the number of processors are increased (Fig 3C).

Personal computers with quad processing cores can run successfully with four workers, which sped up the runtime about 3.5 times with the example images. Thus, even in the absence of a computing core, one can significantly improve the efficiency of the algorithm on a personal computer.

### Distribution of overlapping cell locations

Phase contrast microscopy, which produces a sharp contrast between cells and background, is in general preferable for yeast segmentation and tracking. Yet phase imaging always produces a phase halo around objects [32] that might produce ‘false’ cell boundaries in the context of densely packed cells (Fig 4A). When these ‘false’ boundaries invade the neighboring cells, the segmentation algorithm might assign the same pixels to multiple cells in a way that their segmentations overlap (Fig 4B and 4C, white pixels in Cell Locations).

**Fig 4.**
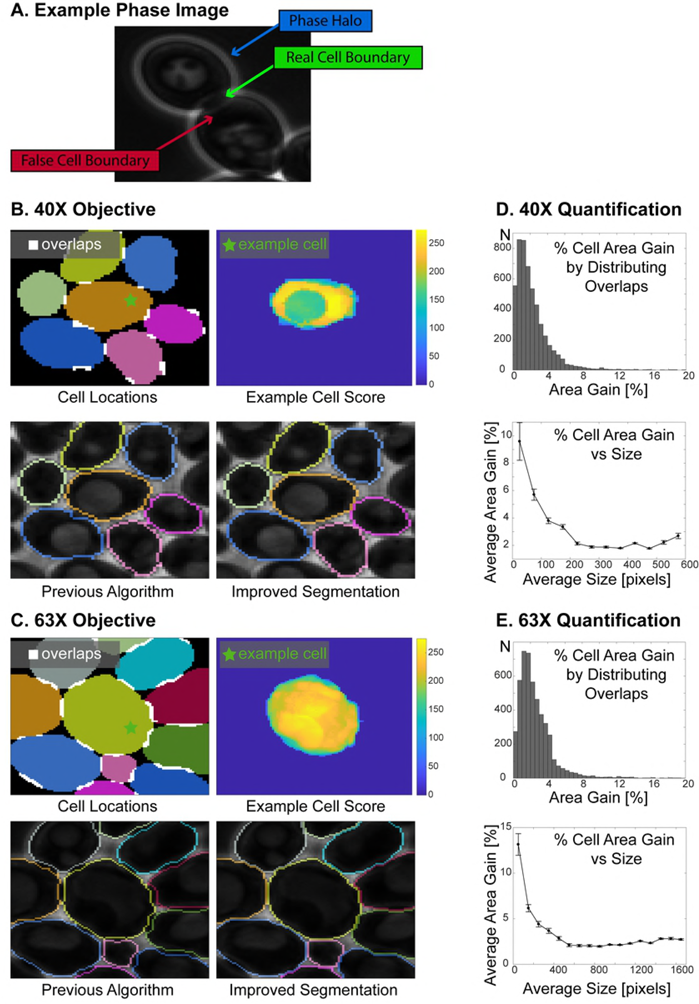
Distribution of overlapping cell locations. **(A)** Example phase image showing two neighboring cells: There is a bright halo (phase halo) around the cells in phase images. When cells are touching, these halos can create a false cell boundary detected by the algorithm. Thus, the algorithm sometimes assigns the same pixels to neighboring cells leading to overlapping cell locations. **(B-C)** Example cells imaged with 40X **(B)** and 63X **(C)** objectives. **Cell Locations**: Overlaps between neighboring cells are highlighted as white areas. Each cell location is represented with a different color. **Example Cell Score**: Each individual cell has a cell score, which carries weights for whether a pixel should belong to the cell. **Previous Algorithm**: Overlapping regions among the cells were excluded from the segmentation in the previous algorithm [30]. **Improved Segmentation**: In the new algorithm such overlapping regions are distributed among the cells based on their scores, which improves the segmentation at the cell boundaries significantly. **(D-E)** Comparison of cell areas with and without distributing the overlapping regions for 40X **(D)** and 63X **(E)** objectives for example cells. Cells imaged over 10 hours (100 time points) were segmented with and without distributing the overlapping pixels. By distributing the overlapping pixels, the majority of cells gained cell area (75% for 40X and 97% for 63X, see Table 2). Percent area gain is calculated by dividing the difference of the cell area with and without distributing the intersections by the area with distributing the intersections and then multiplying the result by 100. Next, the average percent cell area gain versus average size is plotted. To this end, cell sizes are grouped in 50- pixel increments (40X) or in 100-pixel increments (63X). The average size of each group is plotted against the average percent size gain in that group. The error bars show the standard error of the mean. Note that for small cells (buds) area gain percentage is higher than mother cells

**Table 2:**
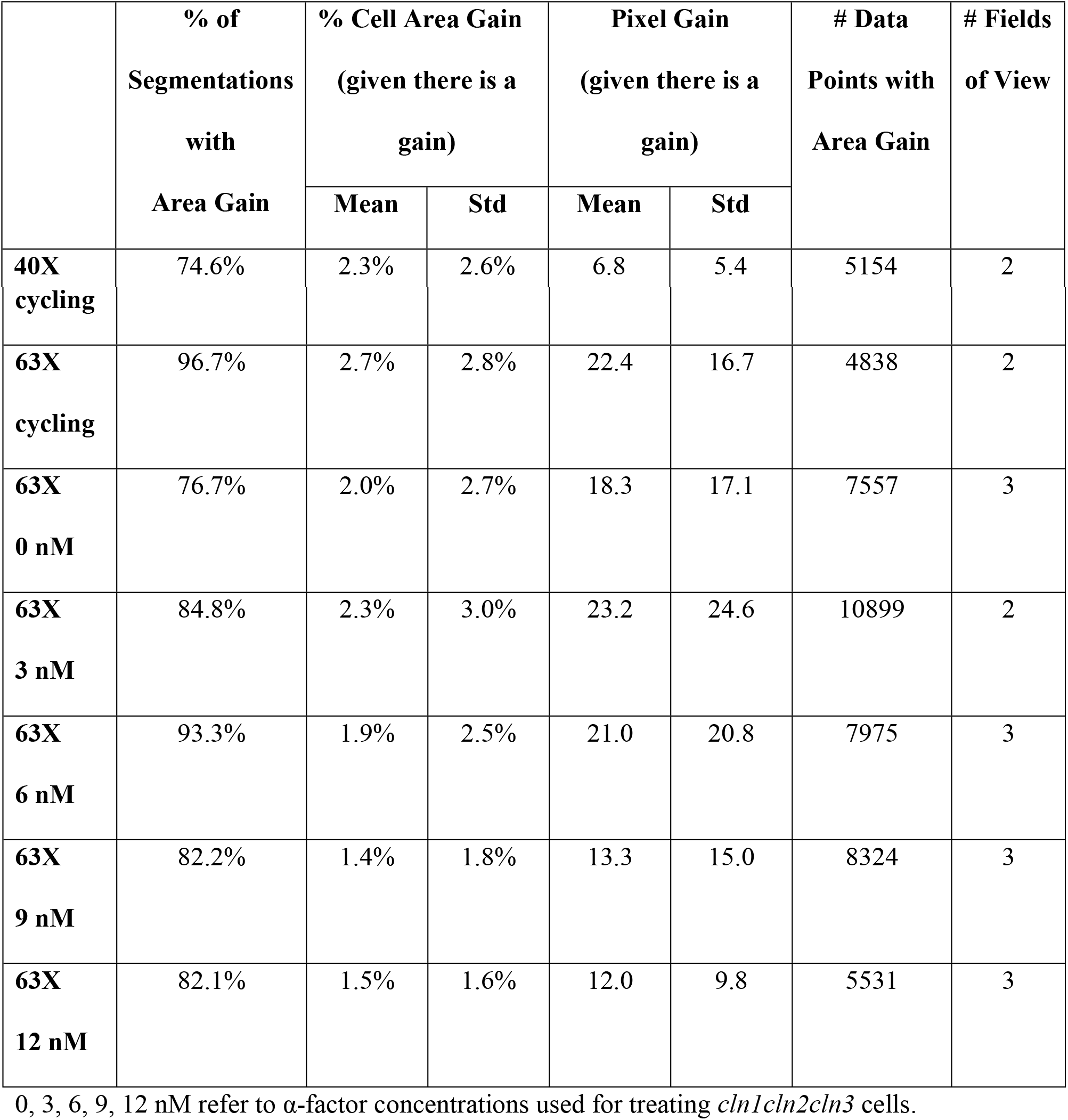
Area Gain by Distribution of Overlapping
Pixels.

In the initial version of our algorithm [30], such overlapping regions were excluded from the segmentation (Fig 4B and 4C, Previous Algorithm). To improve the segmentation accuracy, we developed a method to segment these overlapping areas as well (Fig 4B and 4C, Improved Segmentation). After the cells are segmented individually, the cell locations are compared to detect the overlapping pixels. Next, any such overlapping pixels are distributed based on the scores among cells with overlapping locations. Note that this step is also implemented for automatic seeding (Fig 1D and Fig 2C).

To validate this procedure, we segmented cycling cells imaged for 10 hours (100 time points) with 40X and 63X objectives with or without correction for overlapping cell locations and compared the results (Fig 4 B-E, Movie S1 and S2). Distributing the overlapping regions significantly improved the segmentation as measured by the increase of correctly segmented cell area. Specifically, the vast majority of cells had a non-zero area gain (75%/97% for 40X/63X, Table 2). The cells with an area gain, had increased their area 2.3 ± 2.6% (40X, N_40X_=5154) and 2.7 ± 2.8% (63X, N_63X_=4838) on average. The percent cell area gain is calculated as:

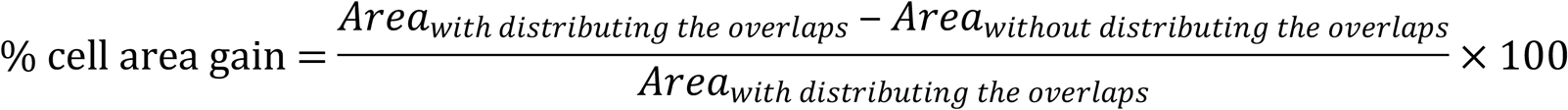

We also tested this correction method for cells with abnormal morphologies. To this end we used a yeast strain that lacks two out of three G1 cyclins (*cln1cln3*) and where the third (*cln2*) was conditionally expressed in our microfluidics-based imaging platform. Specifically, we grew cells for one hour before we arrested the cell cycle and added variable amounts of mating pheromone (0, 3, 6, 9, or 12 nM α-factor) which lead to various yeast morphologies (Fig 5A-E, Movies S3-7) [33, 34]. By distributing the overlapping cell locations, here we noticed again a significant area gain (Table 2). Taken together, this demonstrates that the boundary correction method works and is robust across varying conditions.

**Fig 5.**
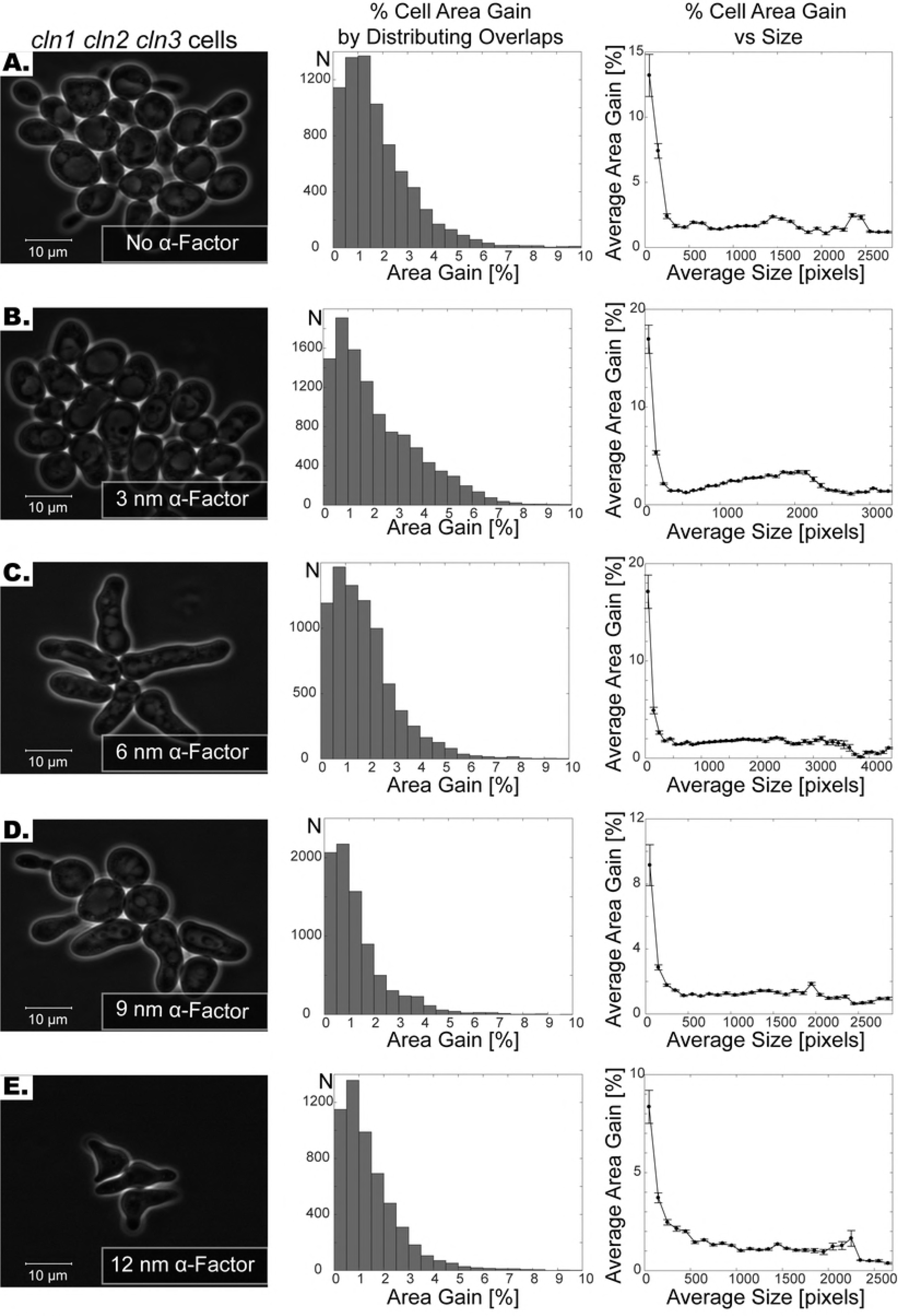
Segmentation of cells subject to varying levels of pheromone treatment. **(A-E)** First column shows the phase images of *cln1 cln2 cln3* cells without α-factor **(A)** and with varying levels of α-factor treatment **(B-E)**. Note that the shapes get progressively more irregular as the concentration of the α-factor increases. Second column shows the histogram of % area gain by distributing the overlapping segmentation regions. Note that histograms are capped at 10%. Third column shows the relationship between size of the cell and the percent cell area gain. The cell sizes are grouped in 100-pixel increments. The average size of each group is plotted against the average percent size gain in that group. The error bars show the standard error of the mean. Note that for small cells area gain percentage is higher than that for larger cells.

### Robustness of segmentation

The ability of a segmentation algorithm to correct an error is a key requirement for correct segmentation over a large number of time points. Otherwise, once an error is made, for example due to an unexpectedly large movement of a cell or a bad focus at one time point, it will linger throughout the segmentation of consecutive time points and errors will accumulate. Our algorithm can correct such errors, since it is robust to perturbations in the seed, i.e. even if there is a segmentation error at one time point, when the algorithm is segmenting the next time point using the previous wrong segmentation as a seed, it can still recover the correct cell boundaries.

To test the robustness of our algorithm to errors in the seed (i.e. segmentation of the previous time point), we picked 340 cells randomly and perturbed their seed by removing 10-90% of the total cell area (Fig 6A). Then, we ran the segmentation algorithm with these perturbed seeds. Over 97% of these cells were fully recovered by the segmentation algorithm (Fig 6B). Out of the 340 cells the algorithm could not recover only 9 cells, which had from 65.5 to 85.9% of their seed removed. On average it took 2.6 ± 2.6 (N=331) time points for the segmentation algorithm to correct segmentation mistakes and the time points required to correct the seed error increased with the severity of the perturbation (Fig 6C). These results demonstrate that our algorithm can correct segmentation mistakes automatically at subsequent time points and, thus, is well suited for long-term imaging.

**Figure 6.**
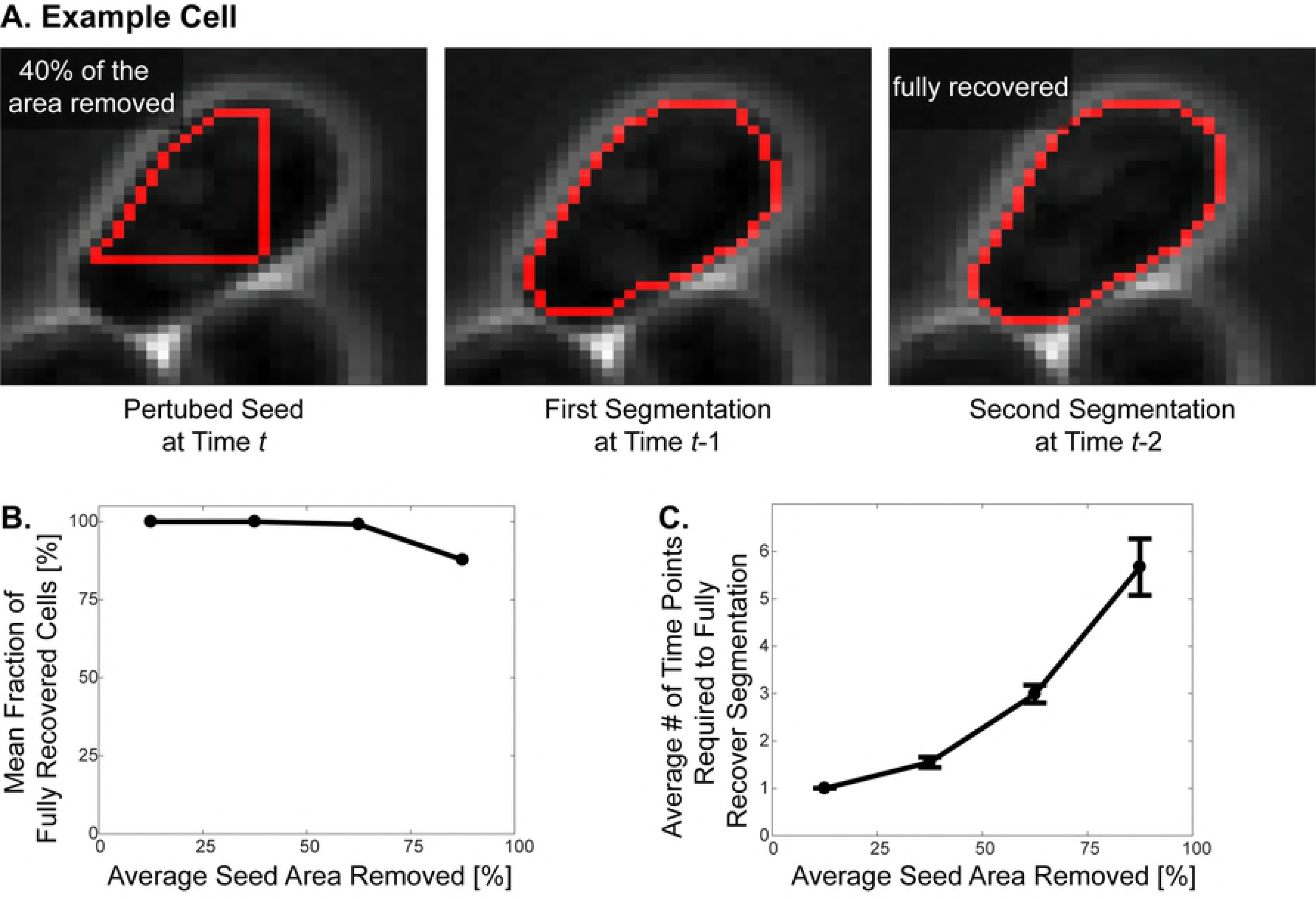
Robustness of the segmentation algorithm to perturbations in the seed. **(A)** Example cell: The seed of the example cell is perturbed by randomly removing 40% of the seed. The algorithm uses this perturbed seed to segment the cell at time *t*-1 and recovers the cell with only minor mistakes. The algorithm fully recovers the cell in two time points. Note that the algorithm segments the cells backwards in time, thus time points are decreasing. **(B)** The seeds of 340 cells were perturbed by randomly removing 10-90% of the seed. The cells are grouped based on the severity of perturbation, i.e. percent seed area removed, in 25% increments. Mean fraction of fully recovered cells are plotted for each group. Note that out of 340 cells, only 9 of them were not recovered by the algorithm. **(C)** The cells are grouped based on the perturbation in 25% increments and the average number of time points required to fully recover the correct cell segmentation is plotted for each group. Number of time points required to fully recover the cells increase with the severity of the seed perturbation. The error bars show standard error of the mean.

Note that the robustness of the algorithm to perturbations is also exploited in the automatic seeding step. Even if the watershed lines produce seeds that are away from the real cell boundary, our algorithm can use those as seed and converge onto the real cell boundaries (Fig 1A). Also, when a cell is over-segmented, i.e. divided into multiple pieces, each piece acts like a perturbed seed and converge onto the correct segmentation. This is why such pieces overlap significantly after running the segmentation subroutine several times (Fig 1D).

## Utilizing fluorescent channels that are not dedicated to segmentation to improve image contrast

A common way to improve segmentation accuracy is to mark cell boundaries by fluorescent dyes or markers [17]. However, such techniques occupy fluorescent channels solely for segmentation, increase the risk of phototoxicity, and/or complicate the experimental setup due to added requirements with respect to cloning (fluorescent proteins) or chemical handling (dyes). It is therefore desirable to limit the number of fluorescent channels dedicated to segmentation if possible.

Nonetheless, *if* any proteins whose localization is at least partially cytoplasmic are fluorescently tagged (dedicated to some biological process of interest), then they can potentially be used to improve the segmentation. Since a large fraction of all proteins exhibit at least partial cytoplasmic localization [35], this is a quite common situation. To take advantage of such cases we developed a method that integrates multi-channel data into the segmentation algorithm. Specifically, this is done by forming a composite image of the phase image (Fig 7A) and the fluorescent channel (Fig 7B), which has high contrast between cell interior and the boundary (Fig 7C).

**Fig 7.**
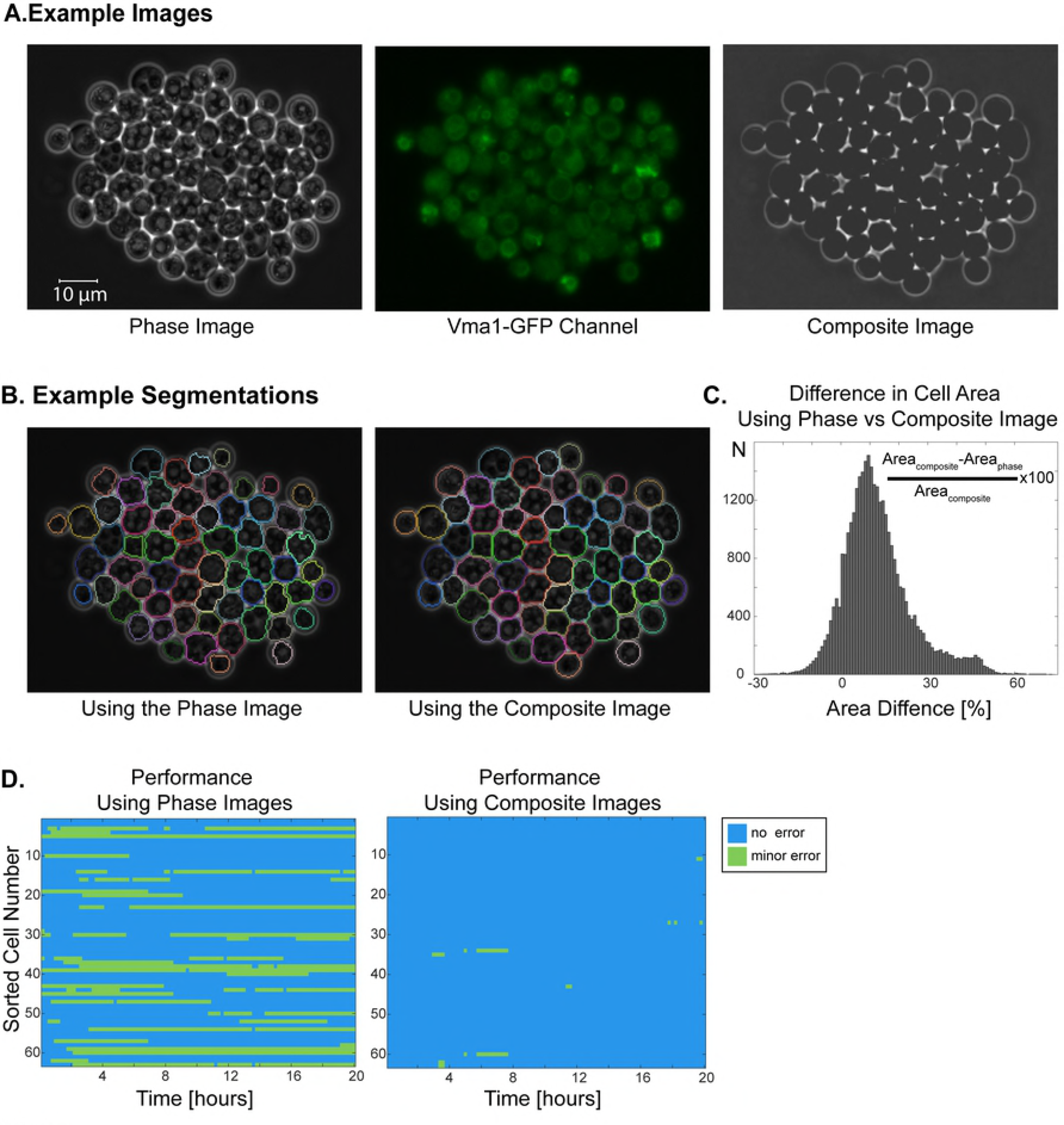
Utilizing a fluorescent channel for improving the segmentation of sporulating cells. **(A)** Example phase image, GFP-channel image and the composite image. In the phase image, spores have very bright patches unlike cycling cells. The composite image is created using the phase and GFP-channel images. Note that Vma1-GFP channel is not dedicated to segmentation. Segmentation results using the phase image and using the composite image. Using the composite image corrects for the slight out of focus phase image and significantly improves the segmentation. **(C-D)** Comparison of segmentations with phase and composite images. Example cells were imaged for 20 hours (100 time points) and segmented with phase or the composite images. **(C)** Out of 32868 cell segmentation events, 89.5% of them have a greater area when the composite image is used for segmentation. **(D)** Comparison of errors in segmentation with phase or composite images. Blue no error, green minor error. Minor errors decreased significantly when using composite images were used for segmentation.

To test this approach, we applied it to yeast cells imaged through the process of spore formation. Such cells, unlike cycling and mating pheromone treated cells, exhibit regions with high phase contrast (white) within the cells (Fig 7A). Moreover, sporulating cells also exhibit morphological changes when the ellipsoidal yeast alters shape to the characteristic tetrahedral ascus shape [36]. Here we used a strain, where the Subunit A of the V1 peripheral membrane domain of the vacuolar ATPase, *VMA1*, is tagged with GFP marking the vacuole boundaries [37]. Note that this biomarker is not dedicated for segmentation; thus, it is a good trial candidate to explore how our method improves segmentation using a biomarker that is not dedicated to segmentation.

We picked two example fields of view, which are segmented over 20 hours (100 time points), amounting to 32868 segmentation events. We segmented these using phase images or composite images. Next, we scored the errors manually and compared the cell areas for each segmentation event that was correctly segmented by both images. We found that 99.3% of the correctly segmented cells had a different cell area and on average they had 12.8 ± 12.0% bigger cell area when composite images are used (Table 3, Fig 7C, Movies S8 and S9). More specifically, we found that 89.5% of the cells have a bigger area when composite image is used for segmentation; 0.7% had the same area, and 9.9% had less cell area. The size distributions of cells segmented using phase and composite images were significantly different (two-sample Kolmogorov-Smirnov test, p<0.001).

**Table 3:** Cell Area with Phase Image and Composite Image.

|  | % of segmentations with area difference | % cell Area Gain (including negatives and zeros) |  | Pixel Gain |  | # Data Points | # Fields of View |
| --- | --- | --- | --- | --- | --- | --- | --- |
|  |  | Mean | Std | Mean | Std |  |  |
| **Sporulating Cells** | 99.3% | 12.8% | 12.0% | 45.4 | 39.8 | 32868 | 2 |

#: Number

In addition, the accuracy of segmentation improved significantly by using composite images. To quantify the accuracy of segmentation, we scored manually the errors in an example field of view, which was segmented with phase images or composite images. A cell is considered accurately segmented if over 95% of its area was segmented correctly. If a segmentation was 90- 95% correct, we labeled it as a minor error. Using composite images, the fraction of correctly segmented cells increased from 75.9% to 99.4% (Table 4, Fig 7D). We found that using the composite image corrects segmentation mistakes that arise due to slightly out of focus phase images.

**Table 4:** Algorithm Performance with Phase Image and the Composite Image.

| Method Used | Phase Only | Composite Image |
| --- | --- | --- |
| # of segmented cells | 63 | 63 |
| # of segmentation events | 6300 | 6300 |
| Fraction of accurate segmentations | 75.9% | 99.4% |
| Fraction of minor segmentation errors | 24.1% | 0.6% |
| Fraction of individual time-series without any errors | 52.4% | 87.3% |
| # Fields of view | 1 | 1 |

#: Number

## Bright-field images

Bright-field images are widely used for live-cell imaging, however they are often low contrast and unevenly illuminated [28]. Thus, it is harder to accurately segment cells using bright-field images.

To test our algorithm on bright-field images, we segmented two example fields of view imaged with bright-field for five hours (100 time points) (Fig 8A, Movie S10). First, we processed the bright-field images to make the cell boundaries more prominent. To this end, we applied top-hat transformation to the complement of the bright-field images (Fig 8B) [38]. We were able to successfully segment bright-field images using our segmentation algorithm (Fig 8C; See section *Overall performance* for quantification of errors).

**Fig 8.**
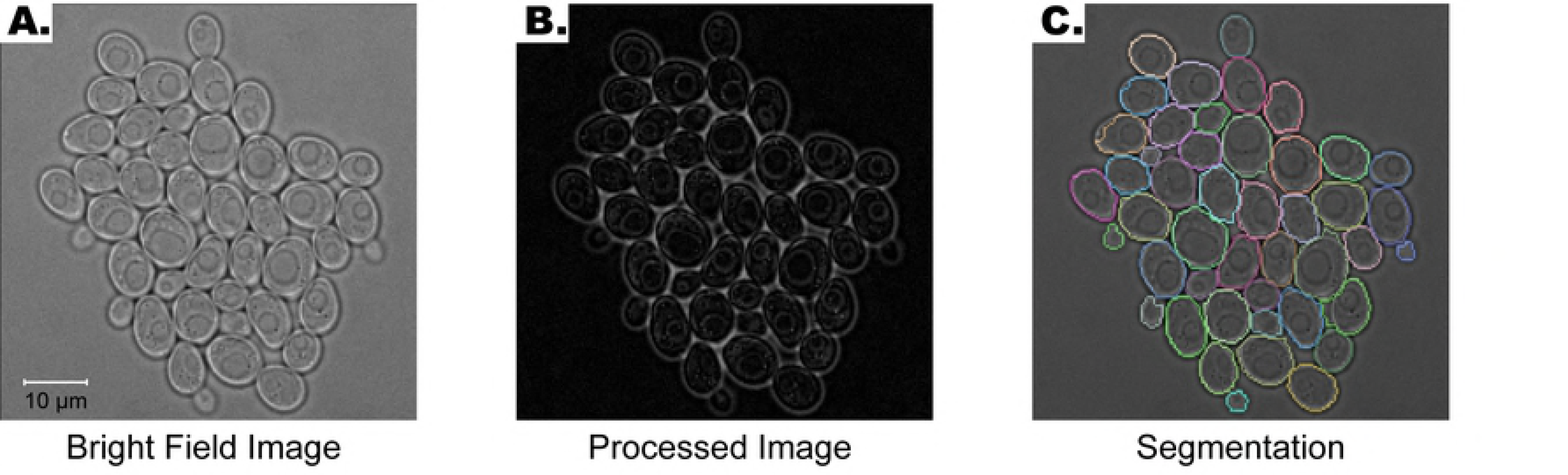
Segmentation of bright-field images. **(A)** Example bright-field image. **(B)** Bright-field image is processed before segmentation by applying a top-hat transform to its complement. **(C)** Segmentation of the image. Each cell boundary is marked with a different color.

## Overall performance

To rigorously test our segmentation algorithm, we segmented 9 different example cases and evaluated our algorithm’s performance. The errors were scored manually. We counted a cell as ‘correctly segmented’ if over 95% of its area was segmented correctly. If the segmentation was 90-95% correct, we labeled it as a minor error. The rest of the errors are called major errors.

The performance of the algorithm is presented in Table 4 and Fig 9. In all example cases at least 92% of the segmentation events were correct. This reached to 99% for some of the example cases. These results demonstrate that our algorithm reaches high accuracy at diverse budding yeast segmentation applications.

**Fig 9.**
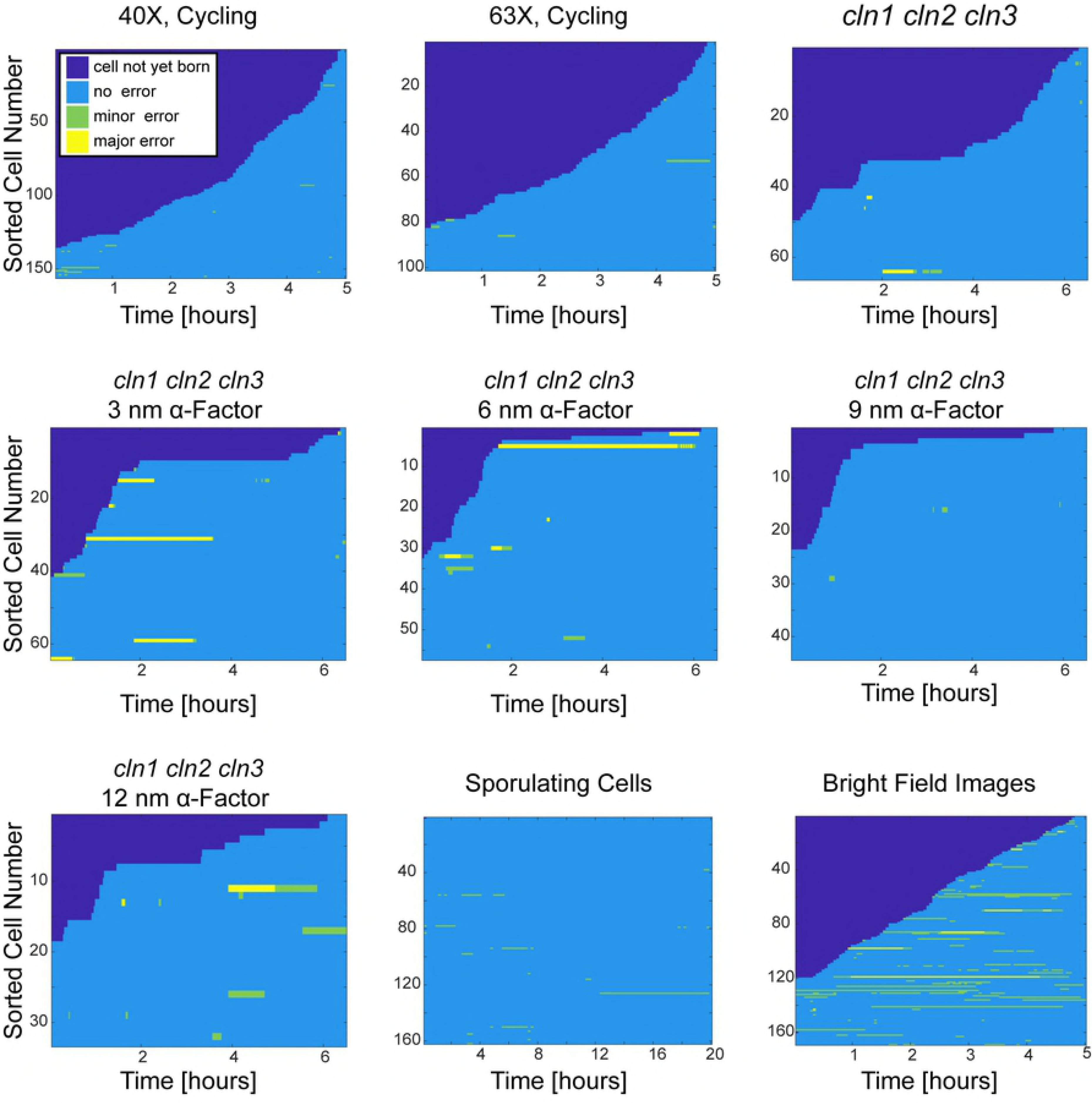
Overall performance of segmentation examples. Sorted cell traces for every example case. Time points where the cell is not yet born are dark blue. Correct segmentations are labeled blue, minor errors green and major segmentation errors yellow. The errors were scored manually. For quantification see Table 4.

**Table 4:**
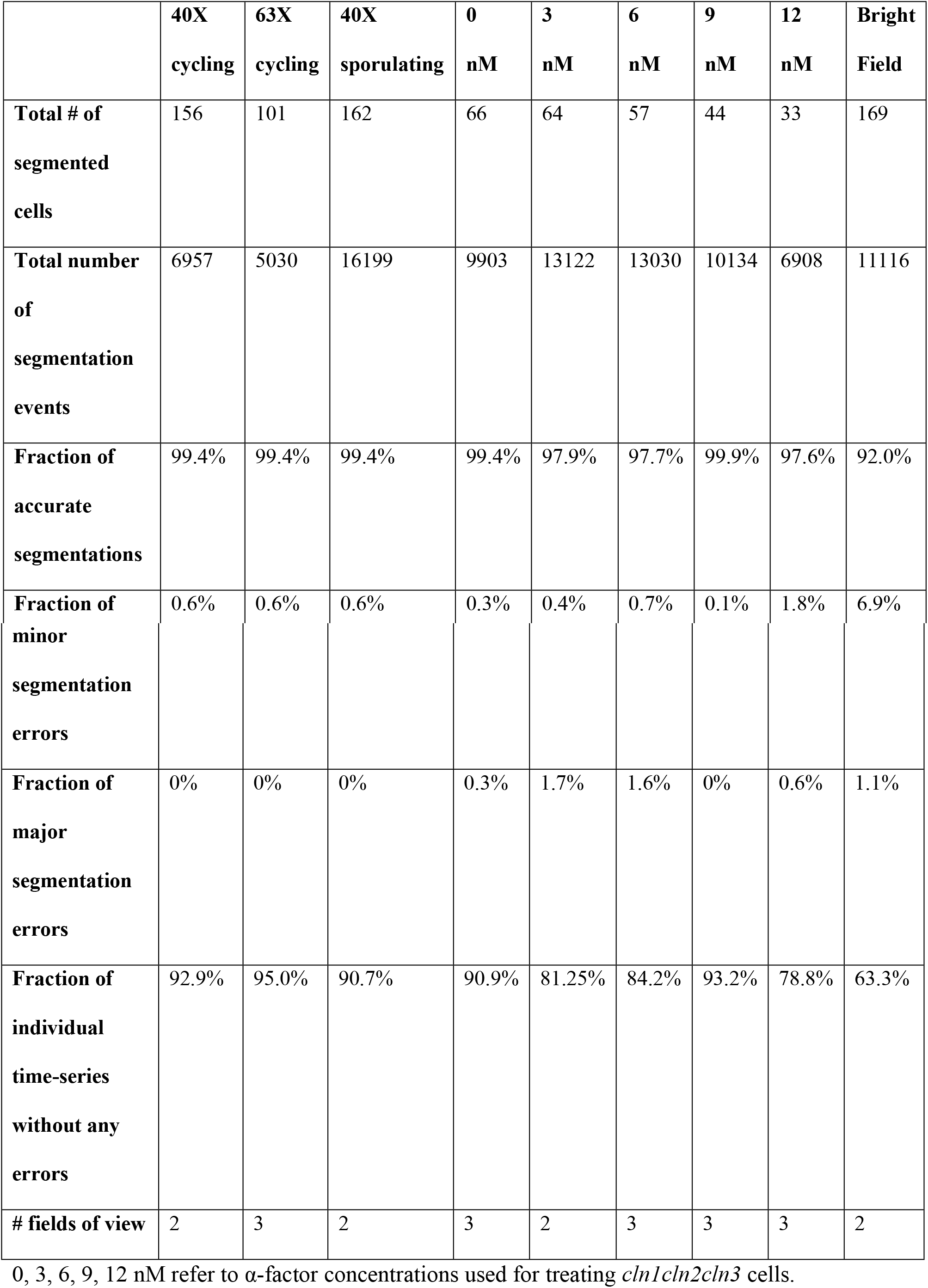
Overall Performance of all Example Cases.

## Discussion

The generation of single cell data from live-cell imaging relies on accurate segmentation and tracking of cells. Once accurate segmentation is achieved, single-cell data can be extracted from a given image time-series [39]. Here we introduce a fully automated and parallelizable algorithm that accurately segments budding yeast cells with arbitrary morphologies imaged through various conditions (phase / bright field) and objectives (40X/63X). This algorithm improves the accuracy and the speed of the previously published one [30] and adapts it to segmentation of different yeast cell morphologies and imaging conditions (Fig 10, Improvements are highlighted in red boxes.). In addition, we developed a novel seeding step, which replaces the semi-automatic seeding of the previous algorithm and enables us to have a fully automatic segmentation algorithm. Since our algorithm can work with no user input, it can be used for large scale single-cell screens.

**Fig 10.**
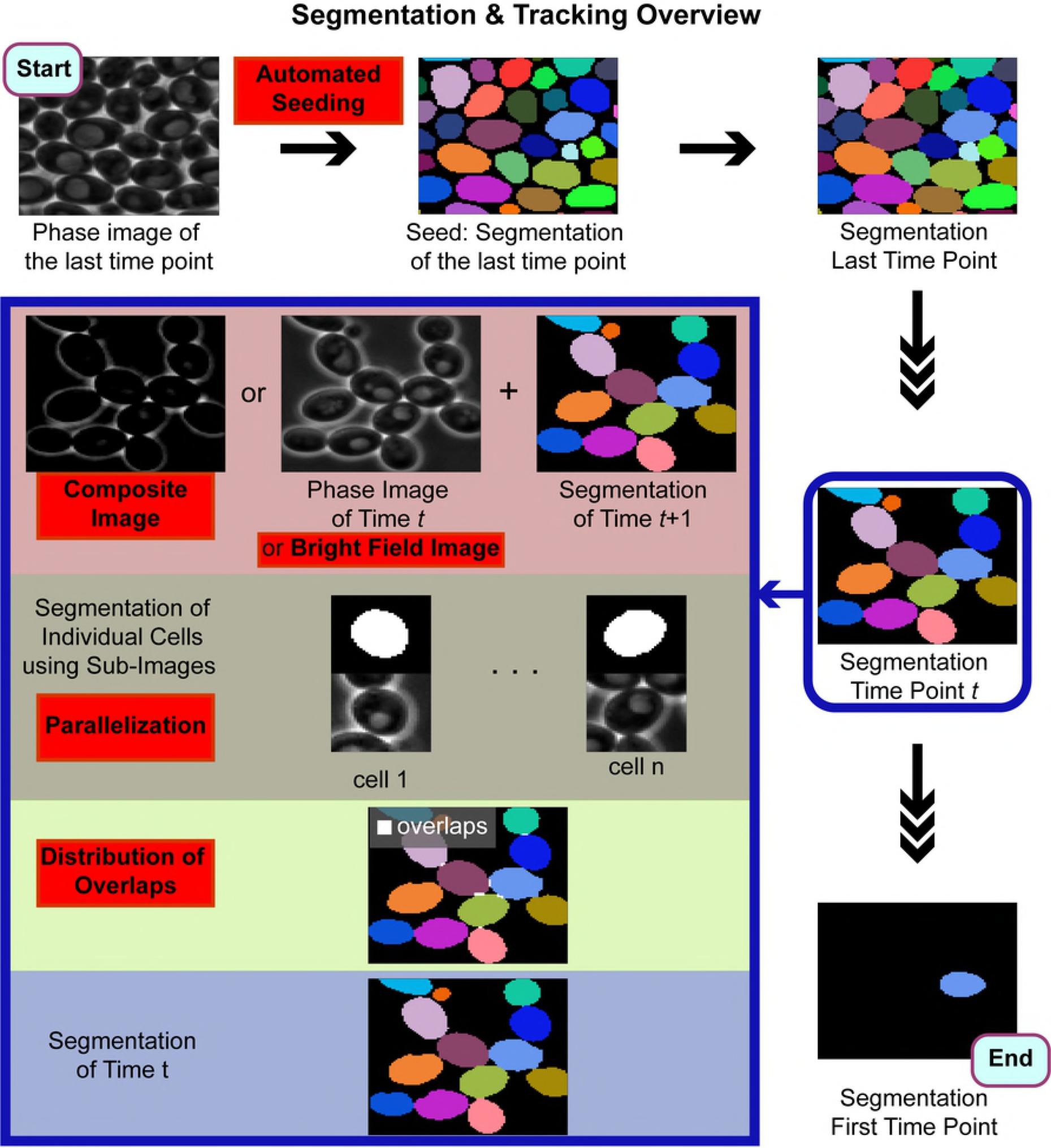
Overview of the segmentation and tracking algorithm. First, the automated seeding step segments the image of the last time point. This seed is fed into the algorithm, which segments the images backwards in time and uses the segmentation of the previous time point as a seed for segmenting the next time point. The segmentation at a given time point is summarized in the blue box. Improvements over the previously published algorithm [30] are highlighted in red boxes.

The algorithm presented here runs significantly faster than the previous algorithm through parallelization. Even in the absence of a computing core, significant time gain can be achieved on a personal computer with two or four processors.

Parallel segmentation of individual cells sometimes leads to assignment of the same pixels to neighboring cells due to false boundaries created by phase halos. Here the algorithm distributes such overlapping regions, which improved the cell area by 1.4-2.8%. The effect of this distribution is more prominent when cells are densely packed. Note that by simply omitting the overlapping pixels the segmentation would still be considered accurate, however, with budding yeast we aim to perfect the segmentation.

Another aim in budding yeast segmentation is to limit the use of fluorescent markers and dyes. Here we show how fluorescent channels that are devoted to a biological process of interest and not to segmentation, can be used to improve the segmentation significantly. The information about the cell location from the fluorescence of the tagged protein and/or autofluorescence of the cells can be incorporated into the phase images by forming composite images using fluorescent channels. In this way, we show a way to utilize existing information about the cell locations in other channels.

One of the strengths of our algorithm is its ability to automatically correct segmentation mistakes at subsequent time points. This property is also exploited in the automatic seeding step to correct mistakes in the seed without user input. Given the versatility and accuracy of our algorithm, we believe that it will improve long-term live cell imaging studies in numerous contexts.

## Materials and methods

### Algorithm Outline

See supplementary material for algorithm outline and the software.

### Media

SCD (1% succinic acid, 0.6% sodium hydroxide, 0.5% ammonium sulfate, 0.17% YNB (yeast nitrogen base without amino acids/ammonium sulfate), 0.113% dropout stock powder (complete amino acid), 2% glucose, YNA [40] (0.25% yeast extract, 2% potassium acetate)

### Cell Culture and Microscopy

The images were taken with a Zeiss Observer Z1 microscope equipped with automated hardware focus, motorized stage, temperature control and an AxioCam HRm Rev 3 camera. We used a Zeiss EC Plan-Neofluar 40X 1.3 oil immersion objective or Zeiss EC Plan-Apochromat 63X 1.4 oil immersion objective. The cells were imaged using a Y04C Cellasic microfluidics device (http://www.cellasic.com/) using 0.6 psi flow rate. Cells were kept at 25 °C. For details of the strains see Table 5.

**Table 5:**
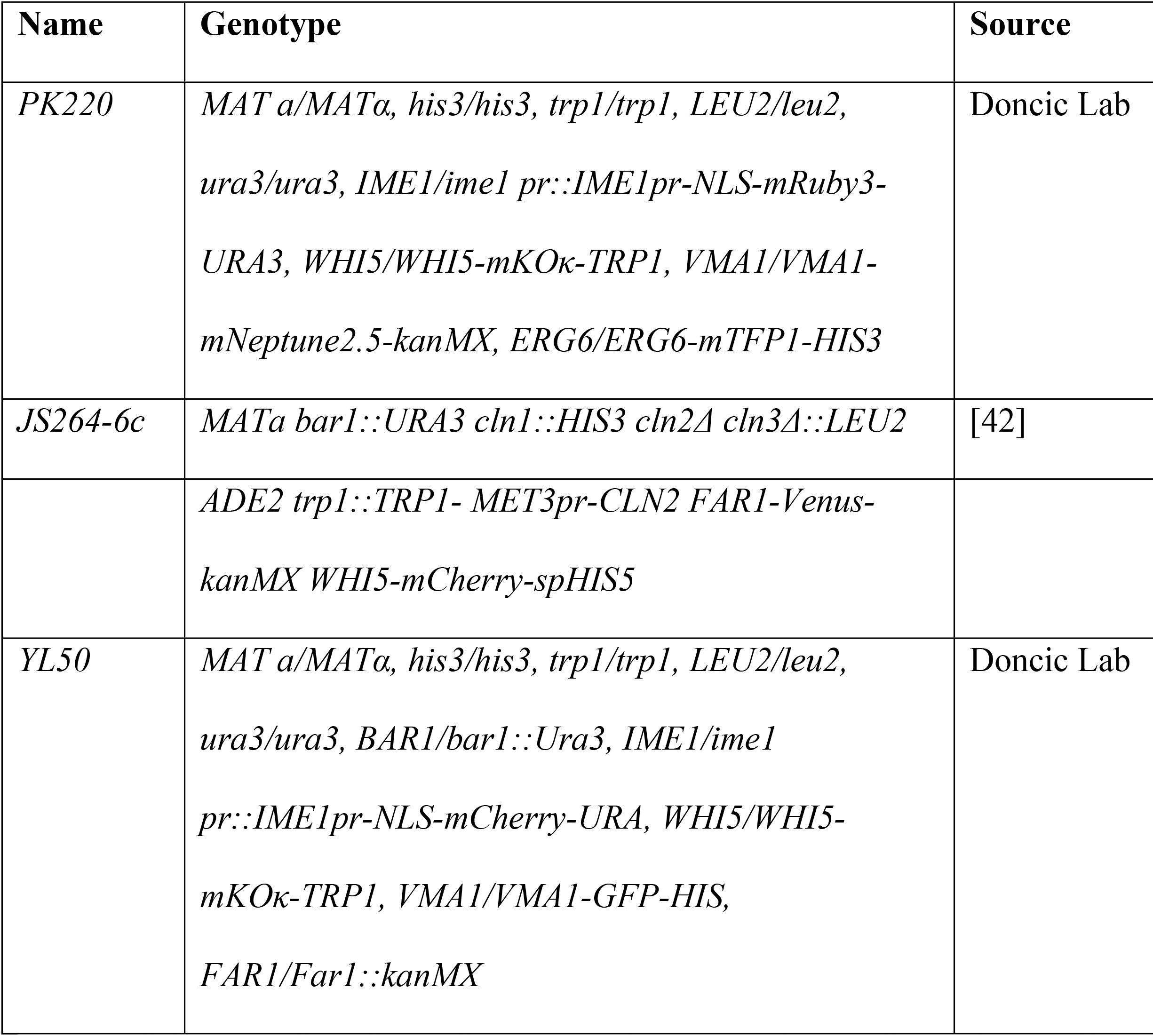
*Saccharomyces Cerevisiae* Strains.

### Cycling Cells

*PK220* cells were imaged in SCD every 3 min with 40X or 63X objective, either with phase contrast or bright field. Exposure times are 40 ms for 40X phase, 80 ms for 63X phase and 20 ms for 40X bright field.

### Sporulating Cells

*YL50* cells were imaged in YNA every 12 min. For details of the sporulation protocol see [41]. Exposure times are 15 ms for phase and 30ms for the GFP channel.

### Pheromone Treated Cells

*JS264-6c* cells received 1h SCD, then they received SCD for 5.5h with mating pheromone (0,3, 6, 9 or 12 nM) and 10X Methionine. Images were taken with 63X objective every 1.5 min.

*JS264-6c* is isogenic with W303 (*leu2-3,112 his3-11,15 ura3-1 trp1-1 can1-1*) and *PK220* and *YL50* are with W303 (ho::LYS2 ura3 leu2::hisG trp1::hisG his3::hisG) except at the loci indicated.

## Acknowledgements

We thank Sandra Schmid and Philippe Roudot for careful reading of the manuscript. We thank Orlando Argüello-Miranda, Yanjie Liu and Piya Kositangool for strains, reagents and help with running the microfluidics experiments. We dedicate this paper to the memory of Andreas Doncic. We hope it will form a part of his scientific legacy.

## Author Contributions

AD and NEW conceived the idea. AD and NEW developed software and analyzed the data. AD and NEW wrote the paper.

## Supporting information

**S1 Movie. Cycling cells imaged with 40X objective.** Cells growing in SCD are imaged every 3 min for 5 hours.

**S2 Movie. Cycling cells imaged with 63X objective.** Cells growing in SCD are imaged every 3 min for 5 hours.

**S3 Movie. *cln1cln2cln3* cells.** Cells growing in SCD are imaged every 1.5 min for 6.5 hours.

**S4 Movie. *cln1cln2cln3* cells exposed to 3 nm α-factor.** The mutant *cln1cln2cln3* cells were grown in SCD for 1 h, and then exposed to 3nM of mating pheromone for 5.5h. The images are taken every 1.5 min.

**S5 Movie. *cln1cln2cln3* cells exposed to 6 nm α-factor.** The mutant *cln1cln2cln3* cells were grown in SCD for 1 h, and then exposed to 6nM of mating pheromone for 5.5h. The images are taken every 1.5 min.

**S6 Movie. *cln1cln2cln3* cells exposed to 9 nm α-factor.** The mutant *cln1cln2cln3* cells were grown in SCD for 1 h, and then exposed to 9nM of mating pheromone for 5.5h. The images are taken every 1.5 min.

**S7 Movie. *cln1cln2cln3* cells exposed to 12 nm α-factor.** The mutant *cln1cln2cln3* cells were grown in SCD for 1 h, and then exposed to 12nM of mating pheromone for 5.5h. The images are taken every 1.5 min.

**S8 Movie. Sporulating cells.** Sporulating cells in YNA are imaged every 12 min for 20 h.

**S9 Movie. Comparison of using composite images vs phase images.** Left is the segmentation of cells using composite images and right are the segmentation of cells using phase images.

**S10 Movie. Bright Field Images.** Cells growing in SCD are imaged every 3 min for 5 hours.

**S11 Text. Tutorial and Algorithm Outline**

**S12 Code and Example Images.**

